# The reovirus μ1 protein contributes to the environmental stability of virions

**DOI:** 10.1101/357343

**Authors:** Anthony J. Snyder, Joseph Che-Yen Wang, Pranav Danthi

## Abstract

The mammalian orthoreovirus (reovirus) outer capsid is composed of 200 μ1-σ3 heterohexamers and a maximum of 12 σ1 trimers. During cell entry, σ3 is degraded by luminal or intracellular proteases to generate a metastable intermediate, called infectious subviral particle (ISVP). Prior to disassembly, σ3 stabilizes the virion by capping μ1. Reovirus fails to establish a productive infection when σ3 degradation is prevented, suggesting proteolytic priming is required for entry. Once uncovered, ISVPs are converted to ISVP*s, which is accompanied by a μ1 rearrangement. Nonetheless, whether σ3 degradation can be bypassed for virions to adopt an altered conformation is undetermined. In this report, we utilized the T1L/T3D M2 reassortant, which encodes a mismatched outer capsid, to further investigate the determinants of reovirus stability. When μ1-σ3 were derived from different strains, virions resembled wild type in structure and protease sensitivity. Using heat as a surrogate for environmental assault, T1L/T3D M2 ISVPs were more susceptible to inactivation than wild type ISVPs. In contrast, virions of each strain were equally stable. Surprisingly, virion associated μ1 rearranged into an ISVP*-like conformation concurrent with loss of infectivity. Despite the presence σ3, a hyperstable variant of μ1 also contributed to heat resistance. The dual layered architecture of reovirus allowed for differential sensitivity to inactivating agents; the inner capsid (core) displayed exceptional resistance to heating. Together, these findings reveal a previously undefined contribution from μ1 in maintaining virion stability.

## Significance

Nonenveloped viruses are exposed to numerous inactivating agents during transmission to a new host. Protein-protein interactions stabilize the particle and protect the viral genome. The mammalian orthoreovirus (reovirus) outer capsid is composed of the μ1 and σ3 proteins. Contacts between neighboring subunits prevent premature disassembly. In this report, we further investigated the determinants of reovirus stability. σ3 controlled heat sensitivity, whereas μ1 enhanced stability under certain conditions. Outer capsid integrity was disrupted at elevated temperatures; virion associated μ1 rearranged into an altered conformation. In contrast, the inner capsid (core) displayed exceptional resistance to heating. Together, these results reveal structural components that differentially contribute to reovirus stability.

## Introduction

Structural integrity is an essential attribute of newly released virions. Perturbations in structure are often accompanied by loss of infectivity (1). Environmental factors can induce irreversible transitions within viral proteins. For example, the capsid of poliovirus adopts an altered conformation in response to elevated temperatures (2, 3), whereas the membrane fusion proteins of influenza virus and Semliki Forest virus are activated by acidic pH (4–6). In each case, the rearrangements that occur *in vitro* resemble those that occur during cell entry. Mammalian orthoreovirus (reovirus) is a useful model system to study many aspects of viral biology (7). Reovirus infectivity is lost upon exposure to heat or chemical agents (8, 9); however, neither the impact on capsid properties nor the mechanism (or basis) of inactivation is fully understood.

Reovirus initiates infection by attaching to proteinaceous receptors (e.g., junctional adhesion molecule A) (10, 11) or serotype specific glycans (e.g., sialic acid or GM2) (12–15). Following internalization by receptor-mediated endocytosis (16–20), virions traffic to Rab7-positive late endosomes in a β1 integrin-, Src kinase-, and microtubule-dependent manner (21–25). Next, acid-dependent cathepsin proteases mediate proteolytic disassembly of the μ1-σ3 outer capsid: σ3 is degraded and μ1 is cleaved into μ1δ and Φ (17, 26–32). The resulting intermediate is called infectious subviral particle (ISVP) (33). Virion-to-ISVP conversion is recapitulated *in vitro* by treating purified virions with chymotrypsin or trypsin (32, 34–37). The metastable ISVP disassembles further to deliver the intact core particle to the host cytoplasm (termed ISVP-to-ISVP* conversion): μ1δ is cleaved into μ1N and δ via autocatalytic activity (38–41). The δ fragment then rearranges into a protease sensitive conformation (38), which is accompanied by the release of μ1N and Φ pore forming peptides (38–46). ISVP-toISVP* conversion is induced *in vitro* using nonspecific factors, such as heat or large, monovalent cations (38, 47, 48), or factors that ISVPs are more likely to encounter during infection, such as μ1N and lipids (49–51).

Reovirus particles exhibit dual layered architecture (33, 41). The inner capsid (core) encapsidates 10 segments of genomic, double-stranded RNA (7). The outer capsid contains 200 μ1-σ3 heterohexamers, which are organized into a T=13 icosahedral lattice (7, 33, 41, 52). The μ1 and σ3 proteins control cell entry and environmental stability. Inhibitors of σ3 degradation block infection (26, 28, 31, 53–55), whereas μ1N cleavage and release are required for penetration of host membranes (42–45). Virion infectivity is susceptible to heat or chemical agents (8, 9). When comparing reovirus strains (e.g., type 1 Lang [T1L] and type 3 Dearing [T3D]), differences in the efficiency of inactivation (inversely related to stability) map to either the M2 gene segment (encodes for μ1) or the S4 gene segment (encodes for σ3) (9). Furthermore, virions are more resistant to thermal inactivation than ISVPs; σ3 confers stability by capping μ1 (41, 48, 52, 54). Mutations in σ3 are correlated with altered sensitivity to heating (53).

We applied structural and biochemical approaches to further investigate the determinants of reovirus stability. T1L and T3D represent the prototype strains for their respective mammalian orthoreovirus serotypes. T1L×T3D reassortants are used extensively to investigate many aspects of the viral replication cycle (7). In this report, we utilized T1L/T3D M2, which encodes the T3D M2 gene and nine genes from T1L. Thus, a mismatched outer capsid is formed between T3D μ1 and T1L σ3. Imperfect interactions may produce subtle changes in structure and/or stability. We demonstrate the following: (i) T1L/T3D M2 virions resemble wild type in structure, protease sensitivity, and thermostability, (ii) virion associated μ1 adopts an altered conformation at elevated temperatures, (iii) a hyperstable variant of μ1 enhances resistance to heating, and (iv) the inner capsid (core) exhibits significant thermostability. These results indicate that μ1 and σ3 contribute to the environmental stability of virions.

## Results

### Recombinant reovirus with a mismatched outer capsid

Reovirus is composed of two concentric, protein shells: the inner capsid (core) and the outer capsid (33, 41). During cell entry, virions are proteolytically disassembled to generate a metastable intermediate, called infectious subviral particle (ISVP) (17, 26–33). The viral and environmental factors that influence ISVP stability have been thoroughly investigated (38, 41, 47–52, 54–67). Notably, heat induces loss of infectivity, which is accompanied by a conformational change within the μ1 outer capsid protein (48). Virions are more resistant to thermal inactivation than ISVPs; the σ3 outer capsid protein confers stability by capping μ1 (41, 48, 52, 54). Nonetheless, whether μ1-σ3 interactions prevent the virion from adopting an altered conformation is undetermined. As such, we utilized the T1L/T3D M2 reassortant to evaluate the effects of a mismatched outer capsid on virion structure and stability. Compared to T1L, the M2 reassortant strain displayed no observable defects in protein composition or protein stoichiometry (Fig. 1A). To rule out gross structural changes, we analyzed virions and ISVPs by dynamic light scattering (DLS). For each virus, we detected a single peak at the expected hydrodynamic diameter (Fig. 1B). Moreover, T1L and T1L/T3D M2 assembled virions that were uniform in size and morphology (Fig. 1C).

**Fig 1.**
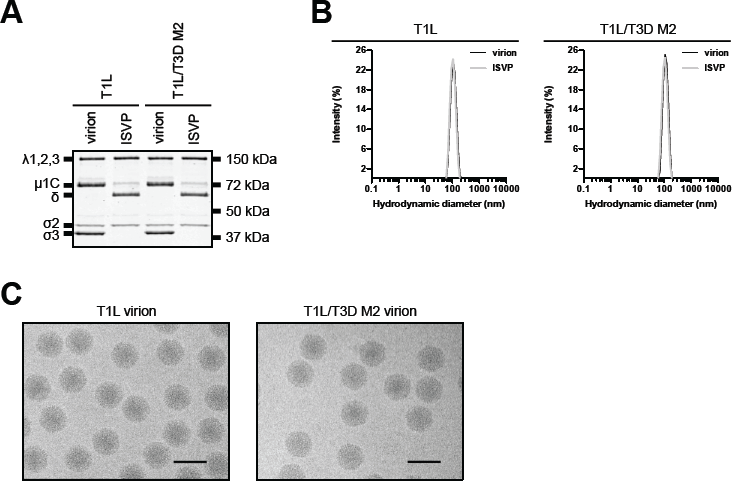
Protein compositions, size distribution profiles, and morphologies of T1L and T1L/T3D M2. (A) Protein compositions. T1L and T1L/T3D M2 virions and ISVPs were analyzed by SDS-PAGE. The gel was Coomassie brilliant blue stained. Migration of reovirus capsid proteins is indicated on the left. μ1 resolves as μ1C, and μ1δ resolves as δ (39). μ1N and Φ are too small to resolve on the gel (n = 3 independent replicates; results from 1 representative experiment are shown). (B) Size distribution profiles.T1L or T1L/T3D M2 virions or ISVPs were analyzed by dynamic light scattering. For each virus, the virion (black) and ISVP (gray) size distribution profiles are overlaid (n = 3 independent replicates; results from 1 representative experiment are shown). (C) Morphologies. T1L or T1L/T3D M2 virions were visualized at nominal 25,000 magnification by cryo-electron microscopy (scale bars = 100 nm; representative images are shown).

### T1L and T1L/T3D M2 virions are structurally equivalent

To further examine the structure of the M2 reassortant strain, we generated cryo-electron microscopy (cryoEM) single-particle reconstructions (Table 1). T1L and T1L/T3D M2 virions displayed common surface features: (i) 600 finger-like projections (represent σ3) that were organized into a T=13 icosahedral lattice and (ii) pentameric turrets (represent λ2) that were present at each of the 5-fold axes of symmetry (Fig. 2A) (33, 41, 52). Minor deviations in density were likely due to differences in resolution. These results were confirmed by comparing central cross sections (Figs. 2B) and by inspecting difference maps (Robem, data not shown). Fitting of the T1L μ1-σ3 heterohexamer verified that T1L and T1L/T3D M2 were similar in structure (Fig. 2C) (52).

**Table 1.** Cryo-electron microscopy single-particle reconstruction statistics.

| Data collection and<br>refinement parameters | Virion |  |
| --- | --- | --- |
|  | T1L | T1L/T3D M2 |
| Micrographs | 205 | 448 |
| Particle number (initial) | 13,415 | 11,751 |
| Particle number (final) | 10,568 | 9,010 |
| Defocus range ($\mu\text{m}$) | 0.5-3.3 | 0.4-4.8 |
| Final resolution ($\text{\AA}$) | 8.2 | 8.8 |
| EMDB<sup>a</sup> accession number | EMD-8916 | EMD-8917 |

<sup>a</sup>Electron Microscopy Data Bank

**Fig 2.**
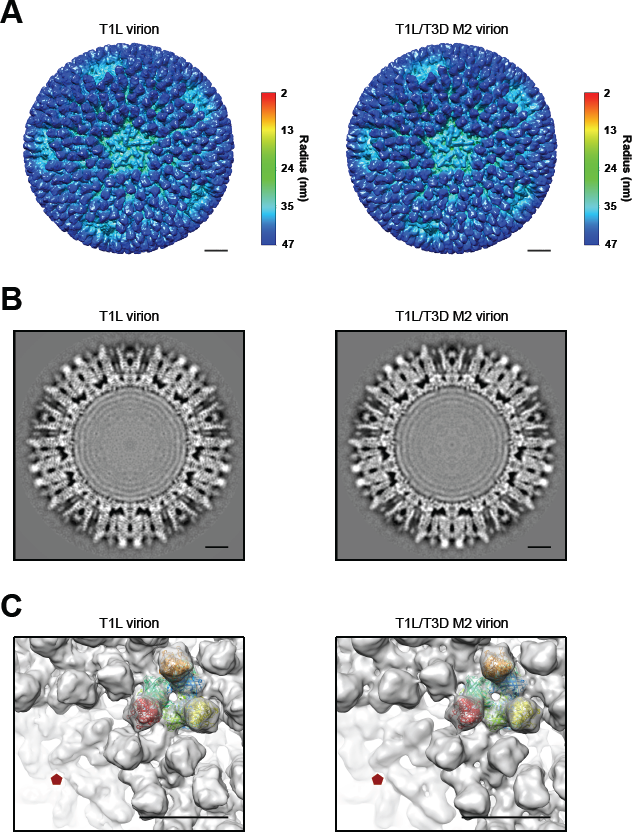
Cryo-electron microscopy single-particle reconstructions of T1L and T1L/T3D M2 virions. (A) Surface shaded representations. Each reconstruction is viewed along an icosahedral fivefold axis. (B) Central cross sections. (C) Fitting of atomic structure. The T1L μ1-σ3 heterohexamer (52) (Protein Data Bank accession number 1JMU) (represented with multicolored ribbons) is docked into each reconstruction (represented with gray densities). Icosahedral fivefold axes are indicated with pentagons (scale bars = 10 nm).

Proteolytic disassembly of the reovirus outer capsid (i.e., virion-to-ISVP conversion) is regulated by a subset of σ3 residues; subtle changes in structure are correlated with altered protease sensitivity (53, 54, 68–70). Mismatched (or imperfect) outer capsid interactions could produce such a change, which may not be discernable at the structural resolutions determined above (Fig. 2). To evaluate differences in σ3 conformation, T1L and T1L/T3D M2 virions were digested *in vitro* with trypsin (TRP) or endoproteinase LysC (EKC) (32, 34, 35, 54, 69). For each virus, TRP catalyzed σ3 degradation within 9 min (Fig. 3A), whereas EKC catalyzed σ3 degradation within 40 min (Fig. 3B). We next tested the sensitivity to intracellular proteases. Reovirus is internalized by receptor-mediated endocytosis (16, 17, 19, 21, 22). Cathepsin B/L then degrade σ3 prior to endosomal escape (17, 26–31). Lysosomotropic weak bases (e.g., ammonium chloride [AC]) block infection by preventing the acidification required for cathepsin activity (31, 71, 72). Thus, the timing of AC escape can be used to monitor the rate of disassembly (26, 28, 31, 53). L cells were adsorbed with T1L or T1L/T3D M2 virions, and viral yield was quantified at 24 h post infection. When indicated, the growth medium was supplemented with AC. Consistent with the above results, each virus bypassed the block to infection at a similar rate (e.g., viral yield of ~1.5 log_10_ units when AC was added at 60 min) (Fig. 3C). Together, the M2 reassortant strain resembles wild type in structure and protease sensitivity.

**Fig 3.**
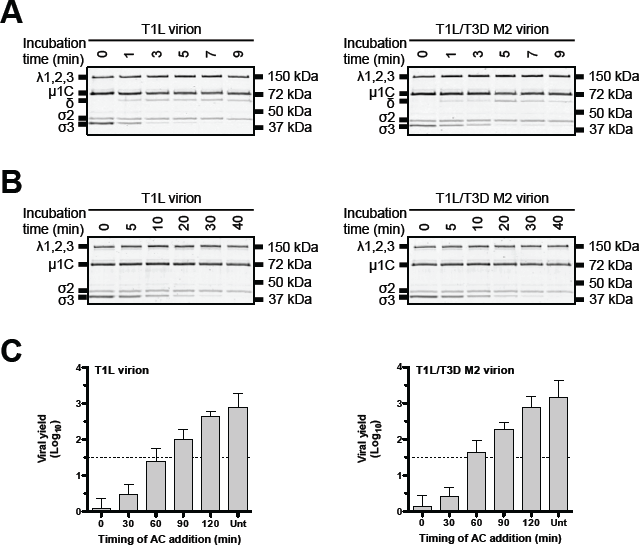
Degradation of the σ3 outer capsid protein by exogenous and intracellular proteases. (A and B) Exogenous proteases. T1L or T1L/T3D M2 virions were incubated in virus storage buffer supplemented with trypsin (A) or endoproteinase LysC (B) for the indicated amounts of time at 8°C (trypsin incubation) or 37°C (endoproteinase LysC incubation). Following digestion, equal particle numbers from each time point were analyzed by SDS-PAGE. The gels were Coomassie brilliant blue stained (n = 3 independent replicates; results from 1 representative experiment are shown). (C) Intracellular proteases. L cell monolayers were infected with T1L or T1L/T3D M2 virions. At the indicated times post infection, the growth medium was supplemented with ammonium chloride. At 24 h post infection, the cells were lysed and viral yield was quantified by plaque assay. Data are presented as means ± SDs (n = 3 independent replicates). AC, ammonium chloride; Unt, untreated.

### T1L ISVPs are more thermostable than T1L/T3D M2 ISVPs

Reovirus ISVPs are formed by the degradation of the σ3 outer capsid protein (17, 26–33). During cell entry, ISVPs undergo an additional conformational change (termed ISVP-to-ISVP* conversion), which culminates in the cleavage and release of μ1 derived pore forming peptides (38–46). When comparing prototype reovirus strains (e.g., T1L and T3D), differences in the efficiency of ISVP-to-ISVP* conversion map to the M2 gene segment (encodes for μ1); T1L μ1 adopts the altered conformation less efficiently than T3D μ1 (38, 39, 48, 60). T1L ISVPs are expected to be more stable than T1L/T3D M2 ISVPs. To confirm these results (and predictions), we conducted 5 min heat inactivation experiments over a range of temperatures. ISVP-to-ISVP* conversion can be triggered *in vitro* using heat. Moreover, ISVPs are infectious, whereas ISVP*s are noninfectious. Thus, thermal inactivation (i.e., loss of infectivity) serves as an indirect measure for ISVP* formation (48). Following incubation at 51°C, T1L ISVPs were reduced in titer by ~4.0 log_10_ units relative to control virus that was incubated at 4°C. In contrast, T1L/T3D M2 ISVPs were reduced in titer by ~2.0 log_10_ units after incubation at 42°C and by ~4.0 log_10_ units after incubation at 48°C (Fig. 4A). Concurrent with ISVP-to-ISVP* conversion, μ1 adopts a protease sensitive conformation (38). This structural rearrangement is assayed *in vitro* by heating ISVPs and determining the susceptibility of the δ fragment (a product of μ1 cleavage) to trypsin digestion (38, 48). Consistent with the thermal inactivation results, the δ fragment in T1L ISVPs became trypsin sensitive at 51°C, whereas the δ fragment in T1L/T3D M2 ISVPs became trypsin sensitive at 42°C (Fig. 4B).

**Fig 4.**
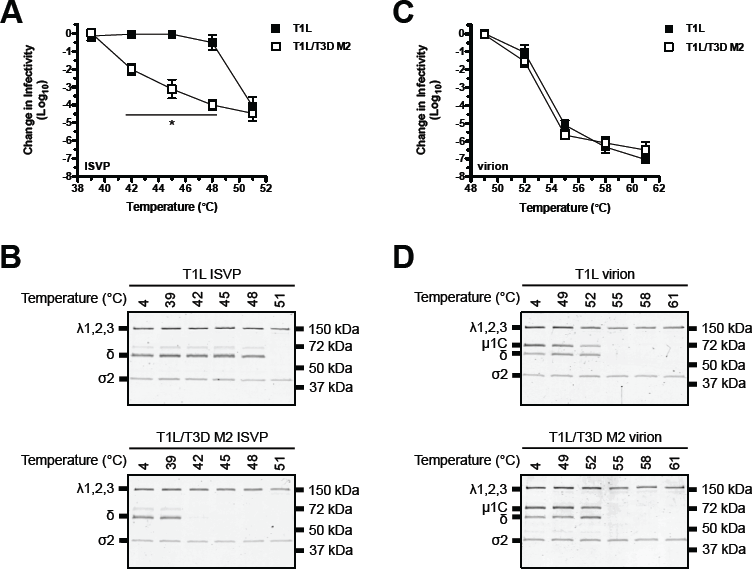
Thermostability of T1L and T1L/T3D M2. (A and C) Thermal inactivation. T1L or T1L/T3D M2 ISVPs (A) or virions (C) were incubated in virus storage for 5 min at the indicated temperatures. The change in infectivity relative to samples incubated at 4°C was determined by plaque assay. Data are presented as means ± SDs. *, *P* ≤ 0.05 and difference in change in infectivity ≥ 2 log_10_ units (n = 3 independent replicates). (B and D) Heat induced conformational changes. T1L or T1L/T3D M2 ISVPs (B) or virions (D) were incubated in virus storage buffer for 5 min at the indicated temperatures. Each reaction was then treated with trypsin for 30 min on ice. Following digestion, equal particle numbers from each reaction were analyzed by SDS-PAGE. The gels were Coomassie brilliant blue stained (n = 3 independent replicates; results from 1 representative experiment are shown).

### T1L and T1L/T3D M2 virions display equivalent thermostabilities

The S4 gene segment (encodes for σ3) controls virion thermostability (9, 48, 54). If σ3 activity (absent in ISVP) is modulated by its interaction with μ1, T1L and T1L/T3D M2 virions might exhibit differential sensitivity to heating. To test this idea, we conducted heat inactivation experiments. Following incubation at 55°C, each strain was reduced in titer by ~5.5 log_10_ units (Fig. 4C). Surprisingly, virion associated μ1 rearranged into an ISVP*-like (i.e., protease sensitive) conformation concurrent with loss of infectivity (Fig. 4D, see 55°C lanes). Of note, σ3 was absent from gels and μ1 migrated as uncleaved μ1C and cleaved δ. Trypsin, which was used to probe for protease sensitivity, degrades σ3 and cleaves at the μ1 δ-Φ junction (32, 34, 35). These results demonstrate that a mismatched outer capsid does not affect thermostability. Moreover, thermal inactivation of virions is correlated with μ1 rearrangements.

### Thermostability of T1L/T3D M2 virions is enhanced by a single μ1 mutation

Virion-to-ISVP conversion exposes particle associated μ1 to solvent (27). Each μ1 monomer, which intertwines with two others to form a trimer, is composed of a jelly roll β barrel domain that connects to a core-adjacent, α-helical pedestal (41, 52). Thus, reovirus ISVPs are stabilized by μ1-mediated intratrimer, intertrimer, and trimer-core interactions (9, 56, 58–62). To determine if μ1 contributes to virion thermostability, we analyzed two variants that exhibit altered efficiency for ISVP-to-ISVP* conversion: T1L/T3D M2 μ1 E89A (reduced thermostability) (61) and T1L/T3D M2 μ1 D371A (increased thermostability) (56, 59). Each virus displayed no observable defects in protein composition, protein stoichiometry, or particle size distribution (Figs. 5A and 5B). Differences in the efficiency of ISVP-to-ISVP* conversion were confirmed by incubating the E89A and D371A variants for 5 min over a range of temperatures. The δ fragment in T1L/T3D M2 ISVPs became trypsin sensitive at 42°C, whereas the δ fragment in T1L/T3D M2 μ1 E89A and T1L/T3D M2 μ1 D371A ISVPs became trypsin sensitive at 33°C and 61°C, respectively (Fig. 5C).

**Fig 5.**
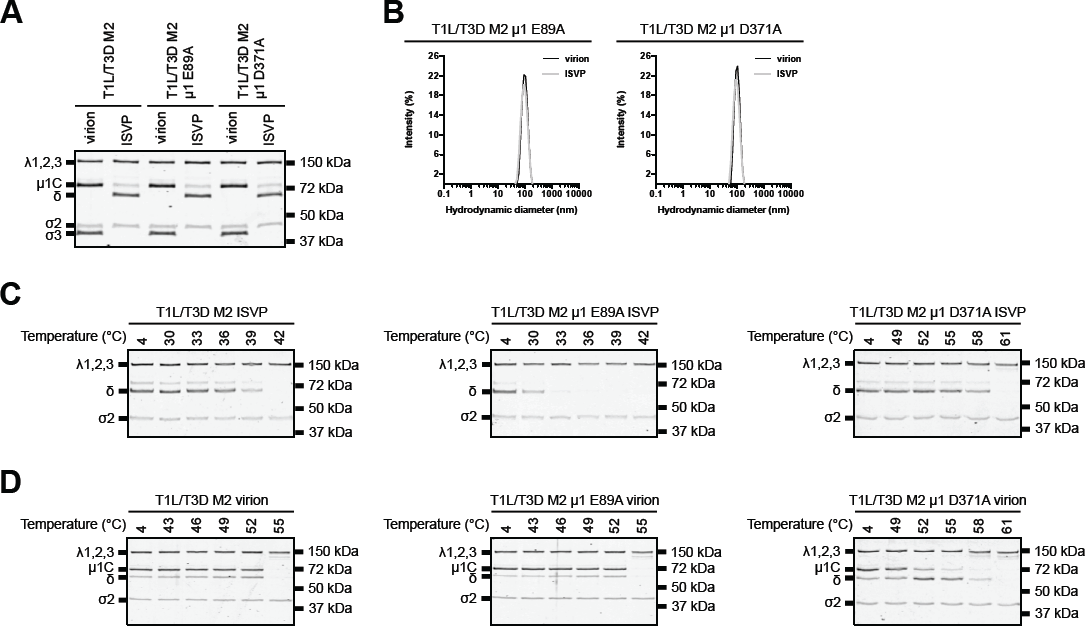
Thermostability of T1L/T3D M2 μ1 variants. (A) Protein compositions. T1L/T3D M2, T1L/T3D M2 μ1 E89A, and T1L/T3D M2 μ1 D371A virions and ISVPS were analyzed by SDS-PAGE. The gel was Coomassie brilliant blue stained. Migration of reovirus capsid proteins is indicated on the left. μ1 resolves as μ1C, and μ1δ resolves as δ (39). μ1N and Φ are too small to resolve on the gel (n = 3 independent replicates; results from 1 representative experiment are shown). (B) Size distribution profiles. T1L/T3D M2 μ1 E89A or T1L/T3D M2 μ1 D371A virions or ISVPs were analyzed by dynamic light scattering. For each virus, the virion (black) and ISVP (gray) size distribution profiles are overlaid (n = 3 independent replicates; results from 1 representative experiment are shown). (C and D) Heat induced conformational changes. T1L/T3D M2, T1L/T3D M2 μ1 E89A, or T1L/T3D M2 μ1 D371A ISVPs (C) or virions (D) were incubated in virus storage buffer for 5 min at the indicated temperatures. Each reaction was then treated with trypsin for 30 min on ice. Following digestion, equal particle numbers from each reaction were analyzed by SDS-PAGE. The gels were Coomassie brilliant blue stained (n = 3 independent replicates; results from 1 representative experiment are shown).

The reovirus outer capsid contains 200 μ1-σ3 heterohexamers (33, 41, 52); σ3 prevents premature disassembly by capping μ1 (9, 48, 54). Nonetheless, virion associated μ1 can adopt an altered conformation at elevated temperatures (Fig. 4). We next tested the idea that virion thermostability is maintained despite μ1 mutations E89A and D371A. For T1L/T3D M2 and T1L/T3D M2 μ1 E89A, μ1 became trypsin sensitive at 55°C. In contrast, T1L/T3D M2 μ1 D371A displayed enhanced resistance to heating; μ1 became trypsin sensitive at 61°C (Fig. 5D). These results indicate the following: (i) σ3 plays a dominant role in stabilizing the virion and (ii) despite the presence of σ3, μ1 can increase thermostability under certain conditions.

### Resistance to chemical agents is controlled by the reovirus outer capsid

To further investigate the factors that regulate stability, chemical agents, such as sodium dodecyl sulfate (SDS) and ethanol, can be used as surrogates to simulate environmental assault (8, 9). The S4 gene segment (encodes for σ3) controls SDS sensitivity (9). Following exposure to 1% SDS, T1L and T1L/T3D M2 virions, which both contain a T1L derived σ3, were reduced in titer by ~2.0 log_10_ units (Fig. 6A). In contrast, ethanol sensitivity is controlled by the M2 gene segment (encodes for μ1); type 1 reoviruses are least sensitive, whereas type 3 reoviruses are most sensitive (9). Following exposure to 33% ethanol, T1L and T1L/T3D M2 virions were reduced in titer by ~1.5 log_10_ units and ~4.0 log_10_ units, respectively (Fig. 6B).

**Fig 6.**
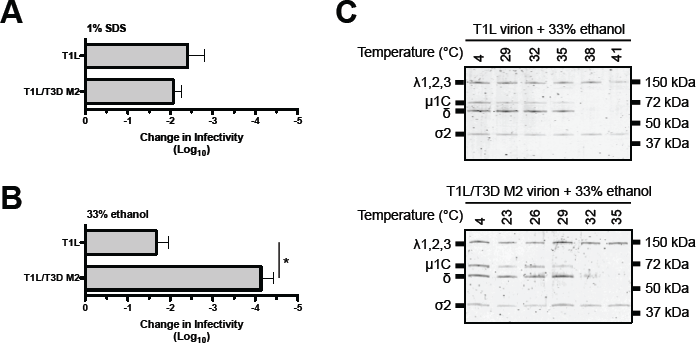
Sensitivity of T1L and T1L/T3D M2 virions to chemical agents. (A and B) Inactivation. T1L or T1L/T3D M2 virions were incubated in virus storage buffer supplemented with 1% sodium dodecyl sulfate (A) or 33% ethanol (B) for 30 min at 42°C (sodium dodecyl sulfate incubation) or 37°C (ethanol incubation). The change in infectivity relative to samples incubated at 4°C was determined by plaque assay. Data are presented as means ± SDs. *, *P* ≤ 0.05 and difference in change in infectivity ≥ 2 log_10_ units (n = 3 independent replicates). (C) Conformational changes. T1L or T1L/T3D M2 virions were incubated in virus storage buffer supplemented with 33% ethanol for 30 at the indicated temperatures. Each reaction was then treated with trypsin for 30 min on ice. Following digestion, equal particle numbers from each reaction were analyzed by SDS-PAGE. The gels were Coomassie brilliant blue stained (n = 3 independent replicates; results from 1 representative experiment are shown).

Single mutations in the δ fragment of T3D μ1 (e.g., A319E, V425F, Q440L, and I442V) (62) or type 3 Abney (T3A) μ1 (e.g., Y466C, K459E, and P497S) (58) confer ethanol resistance. The hyperstable variants perforate membranes less efficiently than wild type (57, 58). As such, we tested whether ethanol inactivation is correlated with μ1 rearrangements. The δ fragment in T1L virions became trypsin sensitive at 38°C, whereas the δ fragment in T1L/T3D M2 virions became trypsin sensitive at 32°C (Fig. 6C). These data provide additional support that μ1 controls virion stability under certain conditions. Moreover, premature disassembly of the outer capsid represents one mechanism (or basis) of inactivation by heat (Figs. 4 and 5) and ethanol (Fig. 6).

### Reovirus core particles exhibit significant thermostability

In our thermostability experiments (Figs. 4 and 5), virions were incubated to temperatures as high as 61°C. Nonetheless, components of the inner capsid (core) (e.g., λ1,2,3 and σ2) remained protease resistant. To confirm this result, T1L and T1L/T3D M2 virions were incubated for 5 min over a range of temperatures, and disassembly was monitored by protease sensitivity. For each virus, the μ1 outer capsid protein became trypsin sensitive at 56°C, whereas the λ1,2,3 and σ2 core proteins became trypsin sensitive at 87°C (Fig. 7B). Interactions between the inner and outer capsids may influence thermostability. To rule out this possibility, core particles derived from T1L and T1L/T3D M2 were purified (Fig. 7A) and then analyzed for heat resistance. Consistent with the above results, the λ1,2,3 and σ2 core proteins became trypsin sensitive at 85°C (Fig. 7C). Of note, we observed an intermediate cleavage fragment when virions or core particles were incubated at 80°C (Figs. 7B and 7C, indicated with an asterisk); peptides were identified that map to λ1 and λ2 (LC/MS-MS, data not shown). Collectively, the dual layered architecture of reovirus allows for differential sensitivity to inactivating agents.

**Fig 7.**
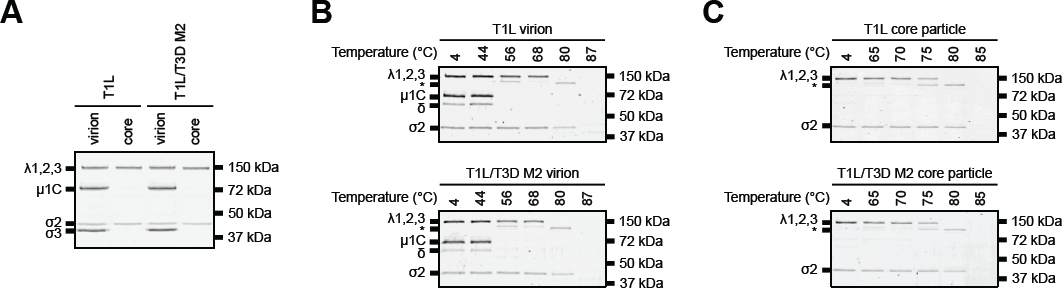
Thermostability of T1L and T1L/T3D M2 core particles. (A) Protein compositions. T1L and T1L/T3D M2 virions and core particles were analyzed by SDS-PAGE. The gel was Coomassie brilliant blue stained. Migration of reovirus capsid proteins is indicated on the left. μ1 resolves as μ1C (39) (n = 3 independent replicates; results from 1 representative experiment are shown). (B and C) Heat induced conformational changes. T1L or T1L/T3D M2 virions (B) or core particles (C) were incubated in virus storage buffer for 5 min at the indicated temperatures. Each reaction was then treated with trypsin for 30 min on ice. Following digestion, equal particle numbers from each reaction were analyzed by SDS-PAGE. The gels were Coomassie brilliant blue stained (n = 3 independent replicates; results from 1 representative experiment are shown).

## Discussion

Reovirus must remain stable during transmission to a new host. Heat or chemical agents can render particles noninfectious (8, 9). Thus, exposure to the environment represents a significant barrier to infection. In this work, we determined the effects of inactivating agents on virion stability and the mechanism (or basis) of inactivation. The T1L/T3D M2 reassortant contains a mismatched outer capsid; imperfect interactions are formed between T3D μ1 and T1L σ3. Reassortment can influence capsid properties (73, 74); however, T1L/T3D M2 resembled wild type in structure and protease sensitivity (Figs. 1-3). During cell entry, virions are converted to a metastable intermediate, called infectious subviral particle (ISVP). Virion-to-ISVP conversion is characterized by σ3 degradation and by μ1 cleavage (17, 26–33). T1L/T3D M2 ISVPs inactivated more efficiently than T1L ISVPs. In contrast, virions of each strain were equally susceptible to elevated temperatures (Fig. 4). Despite the presence of σ3, μ1 also contributed to environmental stability. Following exposure to heat (Fig. 4) or ethanol (Fig. 6), virion associated μ1 rearranged into an altered conformation. Moreover, a hyperstable variant of μ1 enhanced heat resistance (Fig. 5). The dual layered architecture of reovirus allowed for differential sensitivity to inactivating agents. The inner capsid (core) exhibited significant thermostability (Fig. 7). These results reveal a previously undefined contribution from μ1 in maintaining reovirus stability.

Reovirus maintains a balance between structural integrity and conformational flexibility. Virions were less susceptible to heat inactivation than ISVPs (Fig. 4) (48, 54). When comparing strains T1L and T3D, differences in thermostability map to the S4 gene (encodes for σ3); μ1-σ3 interactions confer resistance to inactivating agents (9, 41, 48, 52, 54). Stability is restored by recoating ISVPs with baculovirus-expressed σ3 (54). Moreover, the Y354H mutation in T3D σ3 is correlated with altered sensitivity to heating (53). Nonetheless, a mismatched outer capsid did not affect thermostability. T1L and T1L/T3D M2 virions inactivated at equivalent temperatures (Fig. 4). T1L σ3 and T3D σ3 differ by only 12 residues (97% identical) (multiple sequence alignment, data not shown). The specific interactions that control heat resistance warrant future investigation. In contrast, reovirus ISVPs are stabilized by μ1-mediated intratrimer, intertrimer, and trimer-core interactions (9, 56, 58–62). Notably, the μ1 D371A mutation alters intertrimer contacts (56, 59). This variant enhanced virion and ISVP thermostability (Fig. 5), suggesting μ1 and σ3 cooperate to stabilize the outer capsid.

Reovirus ISVPs undergo a significant conformational change during cell entry. The altered particle is called ISVP* (38). This step is characterized by unwinding and separation of neighboring μ1 trimers (66) and by autocleavage of the μ1 N-terminal fragment (38–41). ISVP-to-ISVP* conversion concludes with the release of pore forming peptides (38–46). Moreover, the μ1 δ fragment adopts a protease sensitive conformation (38). Premature conversion induces loss of infectivity (48); thus, μ1-σ3 interactions are necessary for preserving structural integrity. Nonetheless, virion associated μ1 rearranged into an ISVP*-like (i.e., protease sensitive) conformation at elevated temperatures (Figs. 4). Virion thermostability was enhanced by the μ1 D371A mutation, which is a known regulator of ISVP-to-ISVP* conversion (Fig. 5) (56, 59). Reovirus is also susceptible to chemical agents (8, 9). In particular, ethanol sensitivity is controlled by the M2 gene (encodes for μ1) (9). Mutations in the μ1 δ fragment confer ethanol resistance and reduce membrane penetration efficiency (57, 58, 62). Consistent with the thermostability results, ethanol exposure was correlated with μ1 rearrangements. T1L/T3D M2 was most sensitive, whereas T1L was least sensitive (Fig. 6). Together, heat and ethanol facilitate premature disassembly of the outer capsid. It was undetermined whether μ1-σ3 separate from the particle as a consequence of inactivation. μ1 of recombinantly expressed μ1-σ3 can undergo a rearrangement that resembles ISVP-to-ISVP* conversion. This change requires σ3 degradation and the addition of a triggering agent (e.g., CsCl) (66). Thus, release from the inner capsid (core) is not sufficient for μ1 to adopt a protease sensitive conformation. Preliminary results indicate that the outer capsid was partially or completely lost upon heating (cryo-EM, data not shown). Due to significant aggregation in solution, it was not possible to determine whether μ1-σ3 remained associated (DLS, data not shown).

Nonenveloped, double-stranded RNA viruses are composed of two or three concentric, protein shells (e.g., reovirus, rotavirus, and Bluetongue virus) (75). The outer shell is responsible for cell attachment and membrane penetration. The inner most shell (or core particle) encapsidates the genome and remains intact following entry. Thus, the viral mRNA is transcribed and subsequently released into the host cytoplasm for translation (76, 77). The reovirus core particle exhibited significant thermostability (Fig. 7). The basis for this observation was unclear; however, encapsidation is required for the polymerase to attain an active conformation (78). The hyperstable nature may be necessary for transcription. Interestingly, full activity of the related rotavirus polymerase is also dependent on the interaction with its cognate core particle (79). Nonetheless, rotavirus is significantly less thermostable than reovirus (80). The cause of this difference in heat resistance remains to be identified.

## Materials And Methods

### Cells and viruses

Murine L929 (L) cells were grown at 37°C in Joklik’s minimal essential medium (Lonza) supplemented with 5% fetal bovine serum (Life Technologies), 2 mM L-glutamine (Invitrogen), 100 U/ml penicillin (Invitrogen), 100 μg/ml streptomycin (Invitrogen), and 25 ng/ml amphotericin B (Sigma-Aldrich). All virus strains used in this study were derived from reovirus type 1 Lang (T1L) and reovirus type 3 Dearing (T3D) and were generated by plasmid-based reverse genetics (81, 82). Mutations within the T3D M2 gene were generated by QuikChange site-directed mutagenesis (Agilent Technologies). The μ1 E89A mutation was made using the following primer pair: forward 5’-CGTGCCAAGCGCTTCAGCCTTGGTGCCCTA-3’ and reverse 5’-TAGGGCACCAAGGCTGAAGCGCTTGGCACG-3’. The μ1 D371A mutation was made using the following primer pair: forward 5’-CTCGTGTAGTGAATCTGGCTCAAATCGCTCCGATGCGG-3’ and reverse 5’-CCGCATCGGAGCGATTTGAGCCAGATTCACTACACGAG-3’.

### Virion purification

Recombinant reovirus strains T1L, T1L/T3D M2, T1L/T3D M2 μ1 E89A, and T1L/T3D M2 μ1 D371A were propagated and purified as previously described (81, 82). All strains in the T1L/T3D M2 background contained a wild type or mutated T3D M2 gene in an otherwise T1L background. L cells infected with second passage reovirus stocks were lysed by sonication. Virions were extracted from lysates using Vertrel-XF specialty fluid (Dupont) (83). The extracted particles were layered onto 1.2- to 1.4-g/cm^3^ CsCl step gradients. The gradients were then centrifuged at 187,000 × *g* for 4 h at 4°C. Bands corresponding to purified virions (~1.36 g/cm^3^) (84) were isolated and dialyzed into virus storage buffer (10 mM Tris, pH 7.4, 15 mM MgCl_2_, and 150 mM NaCl). Following dialysis, the particle concentration was determined by measuring the optical density of the purified virion stocks at 260 nm (OD_260_; 1 unit at OD_260_ = 2.1×10^12^ particles/ml) (85). The purification of virions was confirmed by SDS-PAGE and Coomassie brilliant blue (Sigma-Aldrich) staining.

### Generation of infectious subviral particles (ISVPs)

T1L, T1L/T3D M2, T1L/T3D M2 μ1 E89A, or T1L/T3D M2 μ1 D371A virions (2×10^12^ particles/ml) were digested with 200 μg/ml TLCK (*N*α-*p*-tosyl-L-lysine chloromethyl ketone)-treated chymotrypsin (Worthington Biochemical) in a total volume of 100 μl for 20 min at 32°C (36, 37). The reactions were then incubated on ice for 20 min and quenched by the addition of 1 mM phenylmethylsulfonyl fluoride (PMSF) (Sigma-Aldrich). The generation of ISVPs was confirmed by SDS-PAGE and Coomassie brilliant blue (Sigma-Aldrich) staining.

### Generation of core particles

Purified virions of T1L (3.2×10^13^ particles/ml) or T1L/T3D M2 (4.0×10^13^ particles/ml) were digested with 200 μg/ml TLCK-treated chymotrypsin (Worthington Biochemical) in a total volume of 500 μl for 2 h at 37°C (37). The reactions were then incubated on ice for 20 min and quenched by the addition of 1 mM PMSF (Sigma-Aldrich). The digested particles were layered onto 1.3- to 1.5-g/cm^3^ CsCl step gradients. The gradients were centrifuged at 187,000× *g* for 4 h at 4°C. Bands corresponding to purified core particles (~1.44 g/cm^3^) (84) were isolated and dialyzed into virus storage buffer (10 mM Tris, pH 7.4, 15 mM MgCl_2_, and 150 mM NaCl). Following dialysis, the particle concentration was determined by measuring the optical density of the purified core stocks at 260 nm (OD_260_; 1 unit at OD_260_ = 4.4×10^12^ particles/ml) (85). The generation of core particles was confirmed by SDS-PAGE and Coomassie brilliant blue (Sigma-Aldrich) staining.

### Dynamic light scattering (DLS)

T1L, T1L/T3D M2, T1L/T3D M2 μ1 E89A, or T1L/T3D M2 μ1 D371A virions or ISVPs (2×10^12^ particles/ml) were analyzed using a Zetasizer Nano S dynamic light scattering system (Malvern Instruments). All measurements were made at room temperature in a quartz Suprasil cuvette with a 3.00-mm-path length (Hellma Analytics). For each sample, the hydrodynamic diameter was determined by averaging readings across 15 iterations.

### Cryo-electron microscopy and single-particle reconstruction

Purified virions of T1L (9.6×10^13^ particles/ml) or T1L/T3D M2 (4.0×10^13^ particles/ml) were applied to glow discharged, continuous carbon film coated 300-mesh copper grids (Electron Microscopy Sciences). Samples were blotted for 4 s with filter paper and then plunged into liquid ethane using an FEI Vitrobot. All samples were imaged at liquid nitrogen temperatures using a JEOL 3200-FS electron microscope operating at 300 kV. The microscope was equipped with an in-column energy filter. Images were recorded at nominal 25,000 magnification (2.5 Å per pixel) using a Direct Electron DE-20 detector. Each image was collected as 16 individual frames over 1 s. The electron dose per exposure was less than 20 e^-^/Å^2^.

Single-particle reconstructions were processed using EMAN2 and RELION software as previously described (86, 87). Initial models were built *de novo*, and icosahedral symmetry was enforced. Final maps were deposited into the Electron Microscopy Data Bank.

### Modeling

Molecular graphics were created and analyses were performed with the UCSF Chimera package (88).

### Degradation of σ3 by exogenous proteases

T1L or T1L/T3D M2 virions (2×10^12^ particles/ml) were incubated in the presence of 5 μg/ml trypsin (Sigma-Aldrich) or 10 μg/ml endoproteinase LysC (EKC) (New England Biolabs) at 8°C (trypsin digestion) or 37°C (EKC digestion) in a S1000 thermal cycler (Bio-Rad). The starting volume of each reaction was 100 μl in virus storage buffer (10 mM Tris, pH 7.4, 15 mM MgCl_2_, and 150 mM NaCl). At the indicated time points, 10 μl of each reaction was solubilized in reducing SDS sample buffer and boiled for 10 min at 95°C. Equal particle numbers from each time point were analyzed by SDS-PAGE. The gels were Coomassie Brilliant Blue (Sigma-Aldrich) stained and imaged on an Odyssey imaging system (LI-COR).

### Ammonium chloride (AC) escape assay

L cells monolayers in 6-well plates (Greiner Bio-One) were adsorbed with T1L or T1L/T3D M2 virions (10 PFU/cell) for 1 h at 4°C. Following the viral attachment incubation, the monolayers were washed three times with ice-cold PBS and overlaid with 2 ml of Joklik’s minimal essential medium (Lonza) supplemented with 5% fetal bovine serum (Life Technologies), 2 mM L-glutamine (Invitrogen), 100 U/ml penicillin (Invitrogen), 100 μg/ml streptomycin (Invitrogen), and 25 ng/ml amphotericin B (Sigma-Aldrich). The cells were either lysed immediately by two freeze-thaw cycles (input) or incubated at 37°C (start of infection). At the indicated times post infection, the growth medium was supplemented with 20 mM AC (Mallinckrodt Pharmaceuticals). At 24 h post infection, the cells were lysed by two freeze-thaw cycles and the virus titer was determined by plaque assay. The viral yield for each infection condition (i.e. timing of AC addition) (*t*) was calculated using the following formula: log_10_(PFU/ml)*_t_* - log_10_(PFU/ml)input. The titers of the input samples were between 1×10^5^ and 4×10^5^ PFU/ml.

### Thermal inactivation and trypsin sensitivity assays

T1L, T1L/T3D M2, T1L/T3D M2 μ1 E89A, or T1L/T3D M2 μ1 D371A virions, ISVPs, or core particles (2×10^12^ particles/ml) were incubated for 5 min at the indicated temperatures in a S1000 thermal cycler (Bio-Rad). The total volume of each reaction was 30 μl in virus storage buffer (10 mM Tris, pH 7.4, 15 mM MgCl_2_, and 150 mM NaCl). For each reaction condition, an aliquot was also incubated for 5 min at 4°C. Following incubation, 10 μl of each reaction were diluted into 40 μl of ice-cold virus storage buffer (10 mM Tris, pH 7.4, 15 mM MgCl_2_, and 150 mM NaCl) and infectivity was determined by plaque assay. The change in infectivity at a given temperature (*T*) was calculated using the following formula: log_10_(PFU/ml)*_T_* - log_10_(PFU/ml)_4°C_. Under each reaction condition, the titers of the 4°C control samples were between 5×10^9^ and 5×10^10^ PFU/ml. The remaining 20 μl of each reaction were treated with 0.08 mg/ml trypsin (Sigma-Aldrich) for 30 min on ice. Following digestion, equal particle numbers from each reaction were solubilized in reducing SDS sample buffer and analyzed by SDS-PAGE. The gels were Coomassie Brilliant Blue (Sigma-Aldrich) stained and imaged on an Odyssey imaging system (LICOR).

### Sodium dodecyl sulfate (SDS) and ethanol sensitivity assays

T1L or T1L/T3D M2 virions (2×10^12^ particles/ml) were incubated in the presence of 1% SDS (Bio-Rad) or 33% ethanol (Decon Laboratories) for 30 min at 42°C (SDS incubation) or 37°C (ethanol incubation) (Figs. 6A and 6B) or at the indicated temperatures (Fig. 6C) in a S1000 thermal cycler (Bio-Rad). The total volume of each reaction was 30 μl in virus storage buffer (10 mM Tris, pH 7.4, 15 mM MgCl_2_, and 150 mM NaCl). For each reaction condition, an aliquot was also incubated for 30 min at 4°C. Following incubation, 10 μl of each reaction were diluted into 40 μl of ice-cold virus storage buffer (10 mM Tris, pH 7.4, 15 mM MgCl_2_, and 150 mM NaCl), and infectivity was determined by plaque assay. The change in infectivity in the presence of 1% SDS was calculated using the following formula: log_10_(PFU/ml)_42°C_ - log_10_(PFU/ml)_4°C_. The change in infectivity in the presence of 33% ethanol was calculated using the following formula: log_10_(PFU/ml)_37°C_ log_10_(PFU/ml)_4°C_. Under each reaction condition, the titers of the 4°C control samples were between 5×10^9^ and 5×10^10^ PFU/ml. The remaining 20 μl of each reaction were treated with 0.08 mg/ml trypsin (Sigma-Aldrich) for 30 min on ice. Following digestion, equal particle numbers from each reaction were solubilized in reducing SDS sample buffer and analyzed by SDS-PAGE. The gels were Coomassie Brilliant Blue (Sigma-Aldrich) stained and imaged on an Odyssey imaging system (LI-COR).

### Plaque assays

Control or heat-treated virus samples were diluted into PBS supplemented with 2 mM MgCl_2_ (PBS^Mg^). L cell monolayers in 6-well plates (Greiner Bio-One) were infected with 250 μl of diluted virus for 1 h at room temperature. Following the viral attachment incubation, the monolayers were overlaid with 4 ml of serum-free medium 199 (Sigma-Aldrich) supplemented with 1% Bacto Agar (BD Biosciences), 10 μg/ml TLCK-treated chymotrypsin (Worthington Biochemical), 2 mM Lglutamine (Invitrogen), 100 U/ml penicillin (Invitrogen), 100 μg/ml streptomycin (Invitrogen), and 25 ng/ml amphotericin B (Sigma-Aldrich). The infected cells were incubated at 37°C, and plaques were counted at 5 days post infection.

**Statistical analyses.** The reported values represent the mean of three independent, biological replicates. Error bars indicate standard deviation. *P* values were calculated using Student’s *t* test (two-tailed, unequal variance assumed). When comparing experimental outcomes (Figs. 3, 4, and 6), two criteria were used to assign significance: *P* ≤ 0.05 and difference in change in infectivity ≥ 2 log_10_ units.

## Acknowledgements

We thank members of our laboratory and the Indiana University virology community for helpful suggestions. Dynamic light scattering was performed in the Indiana University Physical Biochemistry Instrumentation Facility. Cryo-electron microscopy was performed in the Indiana University Electron Microscopy Center.

Research reported in this publication was supported by the National Institute of Allergy and Infectious Diseases of the National Institutes of Health under award numbers 1R01AI110637 (to P.D.) and F32AI126643 (to A.J.S.) and by Indiana University. The content is solely the responsibility of the authors and does not necessarily represent the official views of the funders.

